## Supplemental Information for "A macroecological perspective on genetic diversity in the human gut microbiome"

### Supporting Information

#### The distribution of allele frequencies under a linearized single-locus model of evolution

##### Deriving the distribution of allele frequencies in the linear purifying selection regime

The derivation of the gamma distribution for a single-locus model of purifying selection in the low frequency limit has previously been described [1]. We begin with a Langevin equation describing evolution on a single-locus with forward and backward mutation ($\mu$, $\upsilon$) and selection ($s$) in a population of $N$ individuals

$$\frac{\partial f}{\partial t} = -sf(1-f) + \mu(1-f) + \upsilon f + \sqrt{\frac{f(1-f)}{N}}\eta(t) \tag{S1}$$

where $\eta(t)$ is a Gaussian noise term. Assuming that $f \ll 1$, the nonlinear selection and back mutation terms drop out and our equation reduces to

$$\frac{\partial f}{\partial t} = -sf + \mu + \sqrt{\frac{f}{N}}\eta(t) \tag{S2}$$

Where I again used the Itô ↔ Fokker-Planck equivalence [2] to formulate a PDE for the probability $p(f,t)$ that an allele has frequency $f$ at time $t$

$$\frac{\partial p(f,t)}{\partial t} = -\frac{\partial}{\partial f}\left[(-sf+\mu)\, p(f,t)\right] + \frac{1}{2N}\frac{\partial^2}{\partial f^2}\left(fp(f,t)\right) \tag{S3}$$

Solving for the stationary distribution, one obtains a gamma distribution of allele frequencies

$$p(f) = \frac{1}{\Gamma\left(2N\mu\right)}\left(2N|s|\right)^{2N\mu}\exp\left[-2N|s|f\right]f^{2N\mu-1} \tag{S4}$$

##### Deriving the distribution of allele frequencies in the linear positive selection regime

We again take the low-frequency limit of the single-locus Langevin, but with $s>0$. This derivation has previously been derived [3, 4], which I rederive below for the convenience of the reader.

$$\frac{\partial f}{\partial t} = sf + \mu + \sqrt{\frac{f}{N}}\eta(t) \tag{S5}$$

This SDE can be solved using the method of characteristics. The solution to this problem has previously been derived [3, 4], which I rederive below for the sake of completeness. To gain a more comprehensive understanding, the unfamiliar reader should refer to the source material. To work with Eq. S5, it will be convenient to define the moment generating function

$$H(z,t) \equiv \left\langle e^{-zf(t)}\right\rangle \tag{S6}$$

which obeys the following PDE

$$\frac{\partial H}{\partial t} = \left[sz - \frac{z^2}{2N}\right]\frac{\partial H}{\partial z} - \mu z H \tag{S7}$$

We then define the log of the moment generating function along the characteristic curve in the absence of mutation ($\mu = 0$) $z^*$ as

$$\Psi(t) \equiv \log\left[H(z^*(t_f - t), t_f - t)\right] \tag{S8}$$

where $tf$ is the current time and $H(z^*(t_f - t), t_f - t)$ is a reverse time function. The function $\Psi(t)$ satisfies the following ordinary differential equation and respective initial conditions, where $\varphi(t) \equiv z^*(t_f - t)$

$$\frac{\mathrm{d}\Psi(t)}{\mathrm{d}t} = \mu\varphi(t) \tag{S9a}$$
$$\Psi(0) = \log\left[H(\varphi(0), t_f)\right] \tag{S9b}$$
$$\Psi(t) = \log\left[H(\varphi(t), 0)\right] = 0 \tag{S9c}$$

which gives us the following function and corresponding moment generating function

$$\Psi(t) = \Psi(0) + \int_0^t \mu\varphi(t^{'})\mathrm{d}t^{'} \tag{S10a}$$
$$H(z,t) = \exp\left[-\mu\int_0^t \varphi(t^{'})\mathrm{d}t^{'}\right] \tag{S10b}$$

At this point, it is necessary to solve $\varphi(t)$, which requires us to look at Eq. S7 for the $\mu = 0$ case. To derive a solution for this case one must identify the family of curves, $z^*$, where $\frac{\mathrm{d}H(z^*,t)}{\mathrm{d}t} = 0$. The line $z^* = 2Ns$ is one characteristic curve. Using the chain rule, one can derive an expression for the ordinary differential equation of $H$

$$\frac{\mathrm{d}H(z^*,t)}{\mathrm{d}t} = \frac{\partial H}{\partial z^*}\frac{\mathrm{d}z^*}{\mathrm{d}t} + \frac{\partial H}{\partial t} = \frac{\partial H}{\partial z^*}\left[\frac{\mathrm{d}z^*}{\mathrm{d}t} + sz^* - \frac{(z^*)^2}{2N}\right] \tag{S11}$$

Solving then the function is equal to zero, one obtains an ordinary differential equation for $z^*$

$$\frac{\mathrm{d}z^*}{\mathrm{d}t} = -sz^* + \frac{(z^*)^2}{2N} \tag{S12}$$

We can then use our definition of $\varphi$ and solve

$$\frac{\mathrm{d}\varphi(t)}{\mathrm{d}t} = s\varphi(t) - \frac{\varphi(t)^2}{2N} \tag{S13a}$$
$$\varphi(t) = \frac{ze^{st}}{1 + \frac{z}{2Ns}\left(e^{st} - 1\right)} \tag{S13b}$$

Finally, one can substitute in the solution for $\varphi(t)$ into Eq.S10a and solve the integral

$$
\begin{aligned}
H(z,t) &= \exp\left[-\mu\int_0^t \frac{ze^{st'}}{1+\frac{z}{2Ns}\left(e^{st'}-1\right)}\mathrm{d}t'\right] && \text{(S14a)}\\
&= \exp\left[-2N\mu\log\left[1+\frac{z}{2Ns}\left(e^{st}-1\right)\right]\right]\Big|_0^t && \text{(S14b)}\\
&= \exp\left[-2N\mu\log\left[1+\frac{z}{2Ns}\left(e^{st}-1\right)\right]\right] && \text{(S14c)}\\
&= \left(1+z\cdot\frac{e^{st}-1}{2Ns}\right)^{-2N\mu} && \text{(S14d)}
\end{aligned}
$$

Which is the moment generating function for the gamma distribution, providing us with a probability distribution of allele frequencies

$$p(f)df = \frac{df}{f_{max}^{2N\mu}\,\Gamma(2N\mu)} f^{2N\mu-1} e^{-f/f_{max}} \tag{S15}$$

where $f_{max} \equiv \frac{e^{st}-1}{2Ns}$ [5]. In the neutral limit $s \to 0$, $f_{max}$ reduces to $t/2N$. This result is corroborated by previous work, where that the maximum size that a neutral mutation can reach over $\sim t$ generations is $\sim t/N$ [5]. This result means that the presence or absence of positive selection alone does not invalidate the gamma distribution as a model for $p(f)$ in the $f \ll 1$ limit.

### 1 Figures

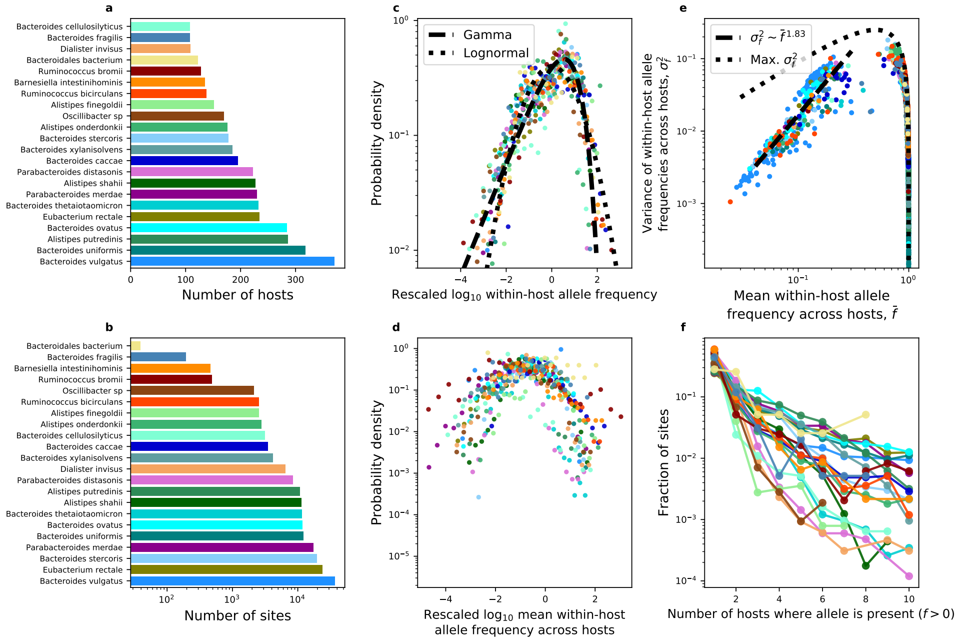


**Figure S1.** Measures of genetic diversity calculated from nonsynonymous sites exhibit similar statistical forms across phylogenetically distant species in the human gut, similar to patterns observed among synonymous sites (Fig. 1).

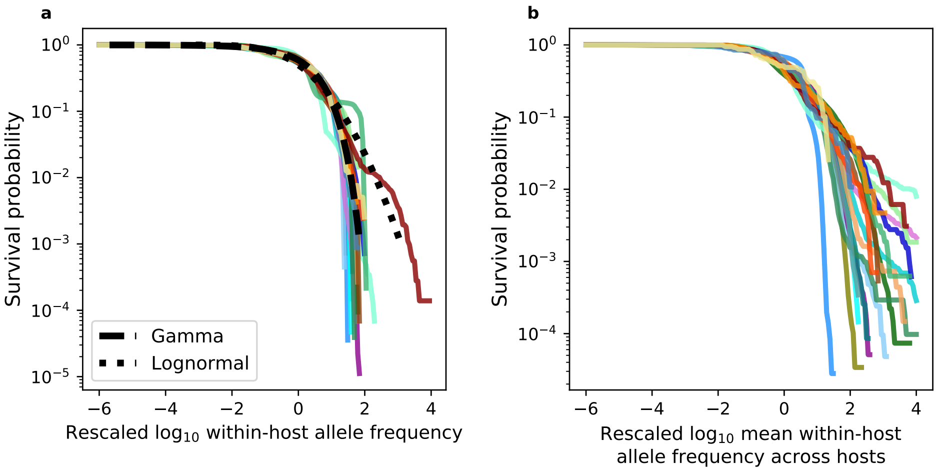


**Figure S2.** Survival forms of rescaled distributions of within-host allele frequencies across hosts and mean frequencies across hosts. Representing the data presented in Fig. 1c,d reveals how distributions of genetic diversity have similar forms across phylogenetically distant species. Each non-black line represents a species. A dashed black line represents the fit of a gamma distribution and dotted black line represents a lognormal.

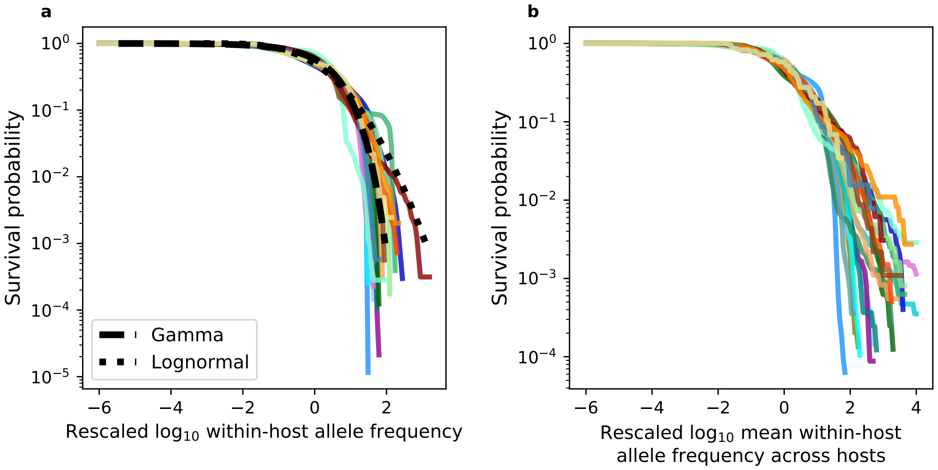


**Figure S3.** The equivalent plot for Fig. S2 for nonsynonymous sites.

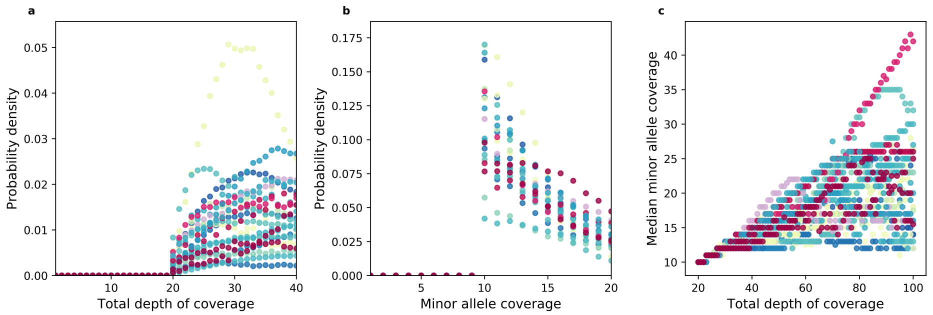


**Figure S4.** The use of the log-likelihood ratio in `MAPGD` introduces a lower bound on the total depth of coverage ($D$) necessary to estimate the frequency of an allele at a given site. **a)** The existence of a lower bound translates to a truncation of the data, where I did not observe any sites with a coverage less than 20 that were processed by `MAPGD`. **b,c)** This truncation means that the depth of coverage of a minor allele ($A$) cannot be less than half the total coverage (e.g., 10).

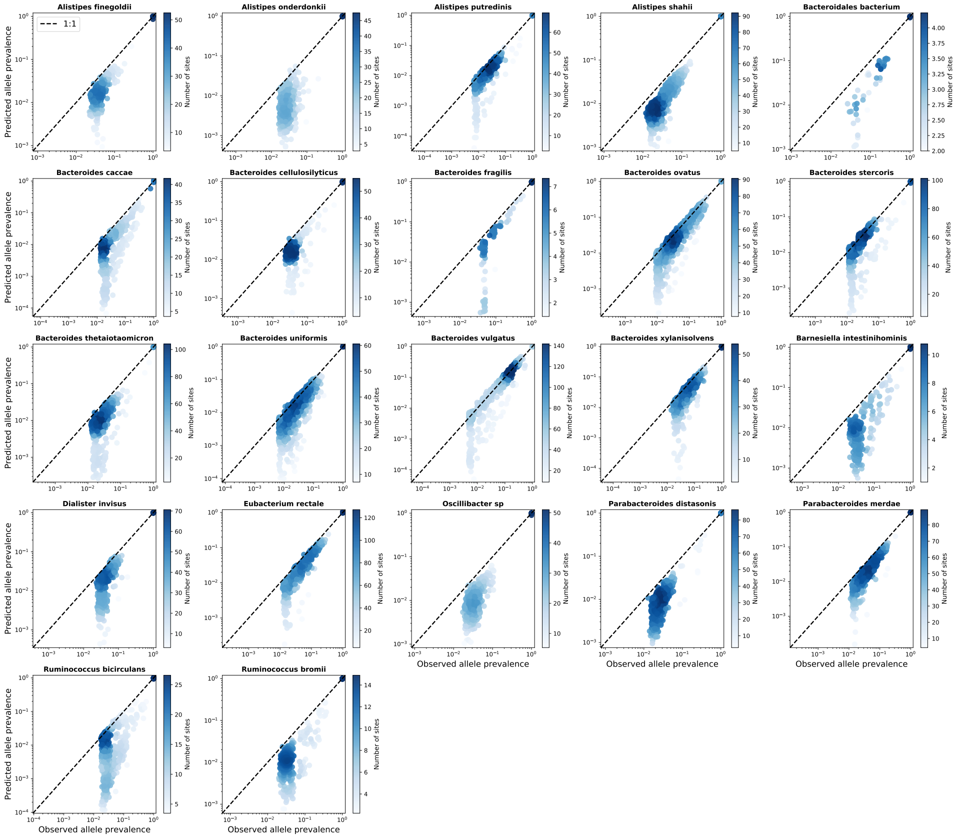


**Figure S5.** A direct comparison between the observed prevalence of all alleles and their corresponding predicted prevalences using the SLM for synonymous sites. A total of 1,000 datapoints were sampled without replacement for each subplot. The color of each datapoint is proportionate to the number of sites.

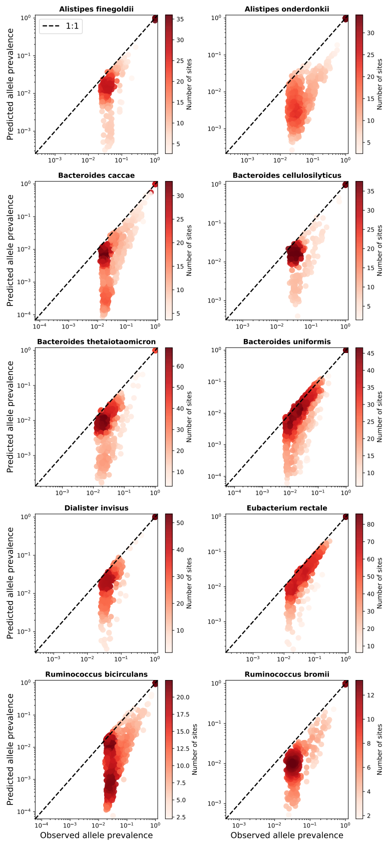


**Figure S6.** A direct comparison between the observed prevalence of all alleles and their corresponding predicted prevalences using the SLM for nonsynonymous sites. A total of 1,000 datapoints were sampled without replacement for each subplot. The color of each datapoint is proportionate to the number of sites.

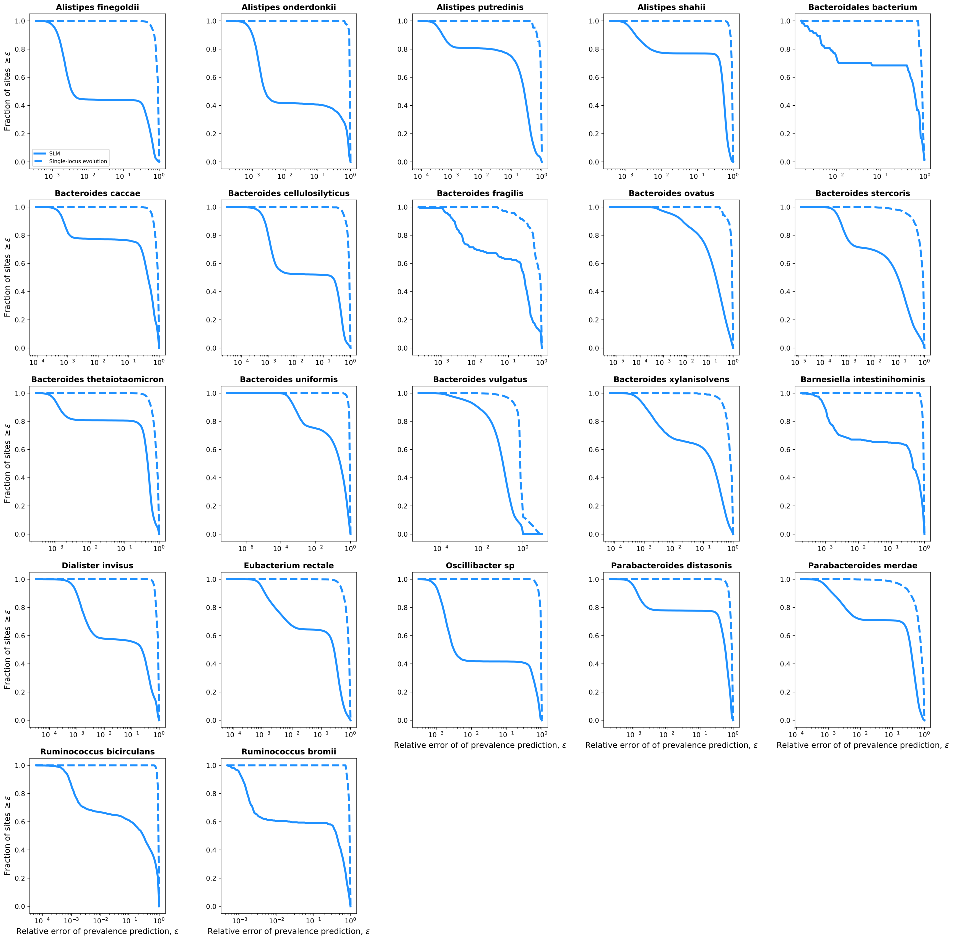


**Figure S7.** By calculating the relative error of all alleles for the SLM we can examine the error distributions. To visually compare the two models, I examined the survival distribution of the relative errors (i.e., the compliment of the empirical cumulative density function). All alleles in this plot are at synonymous sites.

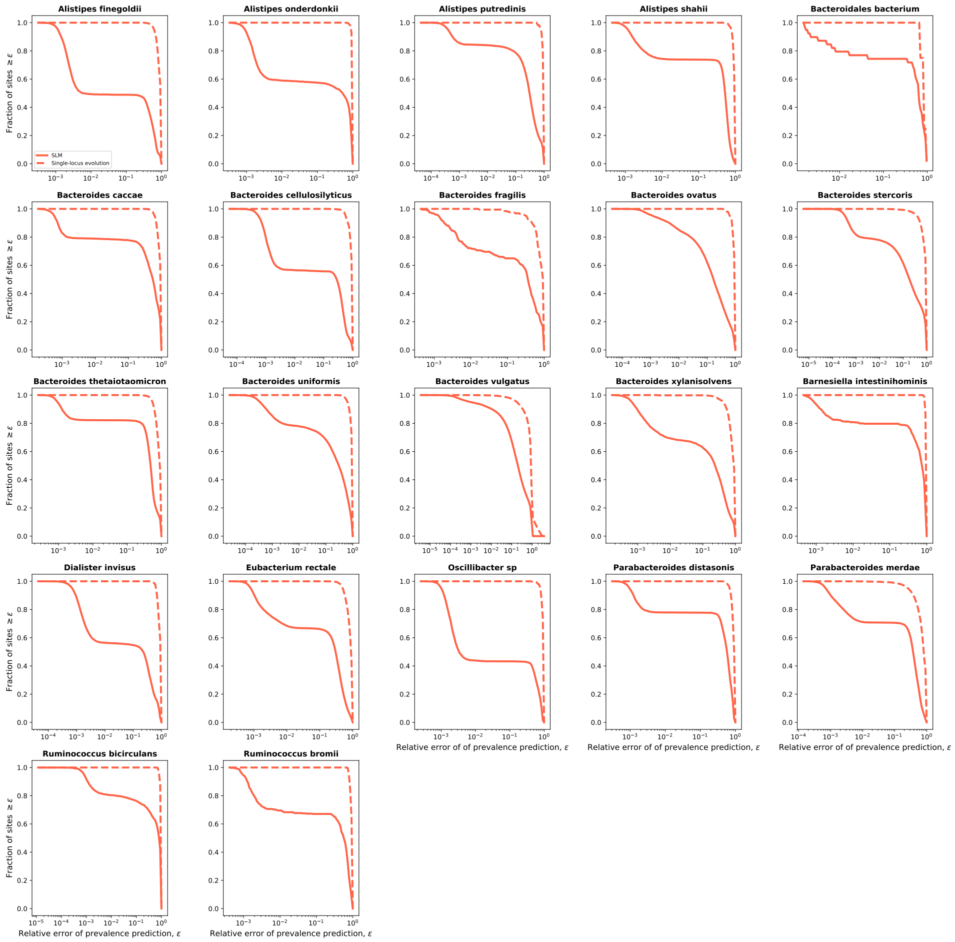

**Figure S8.** By calculating the relative error of all alleles for the SLM we can examine the error distributions. To visually compare the two models, I examined the survival distribution of the relative errors (i.e., the compliment of the empirical cumulative density function). All alleles in this plot are at nonsynonymous sites.

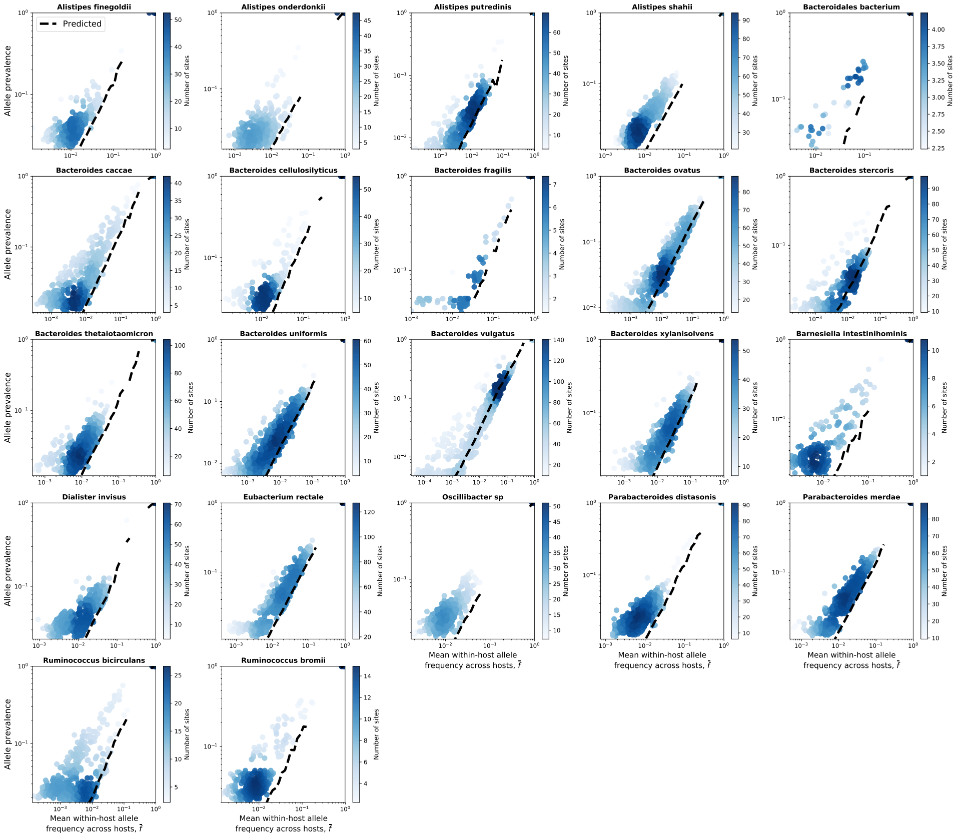


**Figure S9.** The empirical relationship between the mean frequency of an allele ($\bar{f}$) and its prevalence across hosts can be recapitulated by the SLM for synonymous sites. Blue dots represent observed values and the shade of blue is proportional to the density of observations. The black line is the predicted relationship calculated using Eq. 11. A total of 1,000 datapoints were sampled without replacement for each subplot. The color of each datapoint is proportionate to the number of sites.

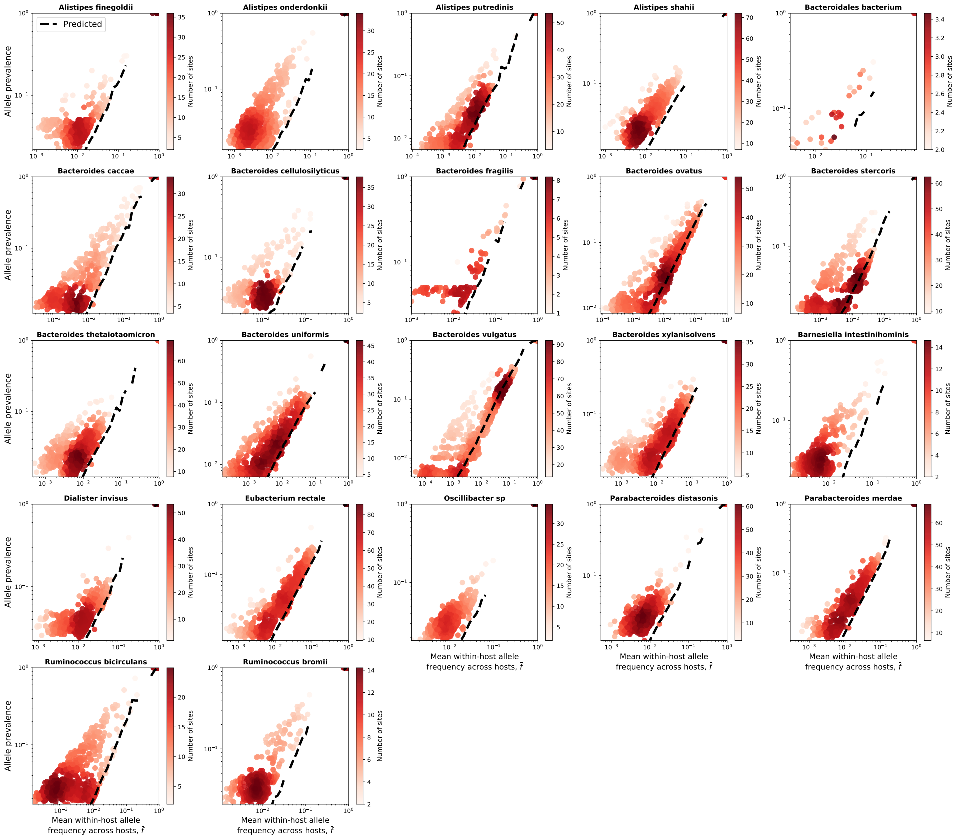


**Figure S10.** The empirical relationship between the mean frequency of a given allele ($\bar{f}$) and its prevalence across hosts can be recapitulated by the SLM for nonsynonymous sites. Blue dots represent observed values and the shade of blue is proportional to the density of observations. The black line is the predicted relationship calculated using Eq. 11. A total of 1,000 datapoints were sampled without replacement for each subplot. The color of each datapoint is proportionate to the number of sites.

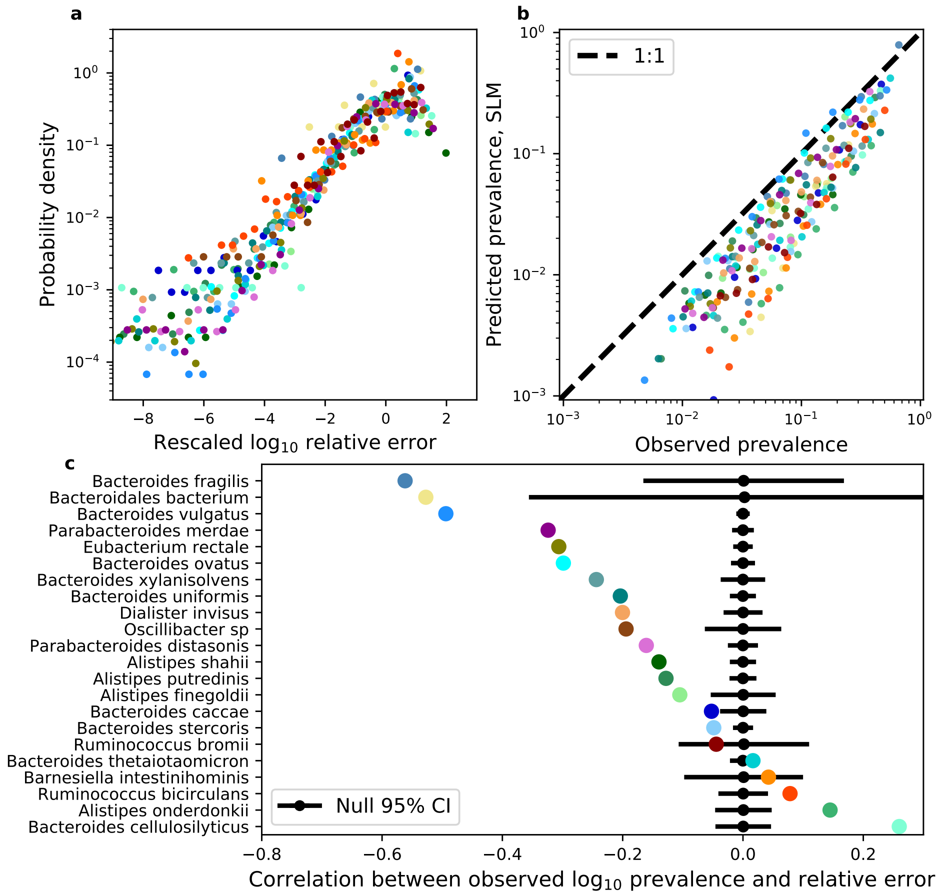


**Figure S11.** The equivalent analyses in Fig. 3 were performed on alleles at nonsynonymous sites. The results of these analyses are qualitatively consistent with those of synonymous sites.

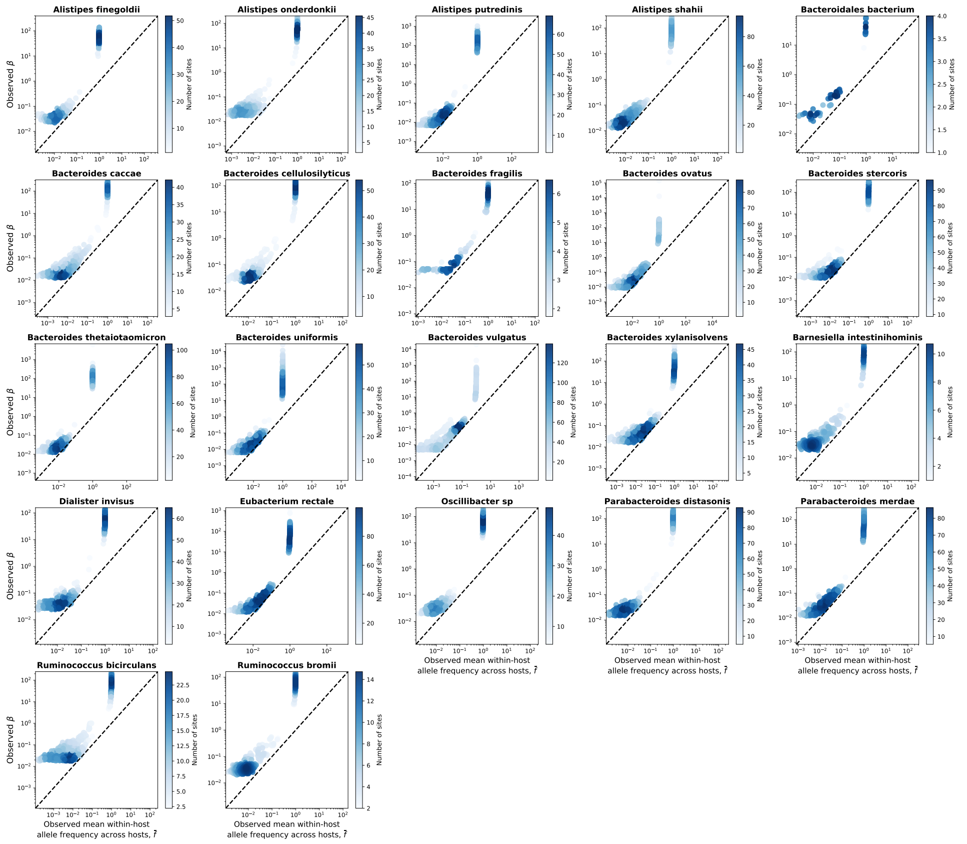


**Figure S12.** The relationship between the empirical estimates of the two parameters of the SLM: the mean allele frequency across hosts ($\bar{f}$) and the squared inverse of the coefficient of variation of frequencies across hosts ($\beta$). Each point is an individual allele. All alleles are on synonymous sites. A total of 1,000 datapoints were sampled without replacement for each subplot. The color of each datapoint is proportionate to the number of sites.

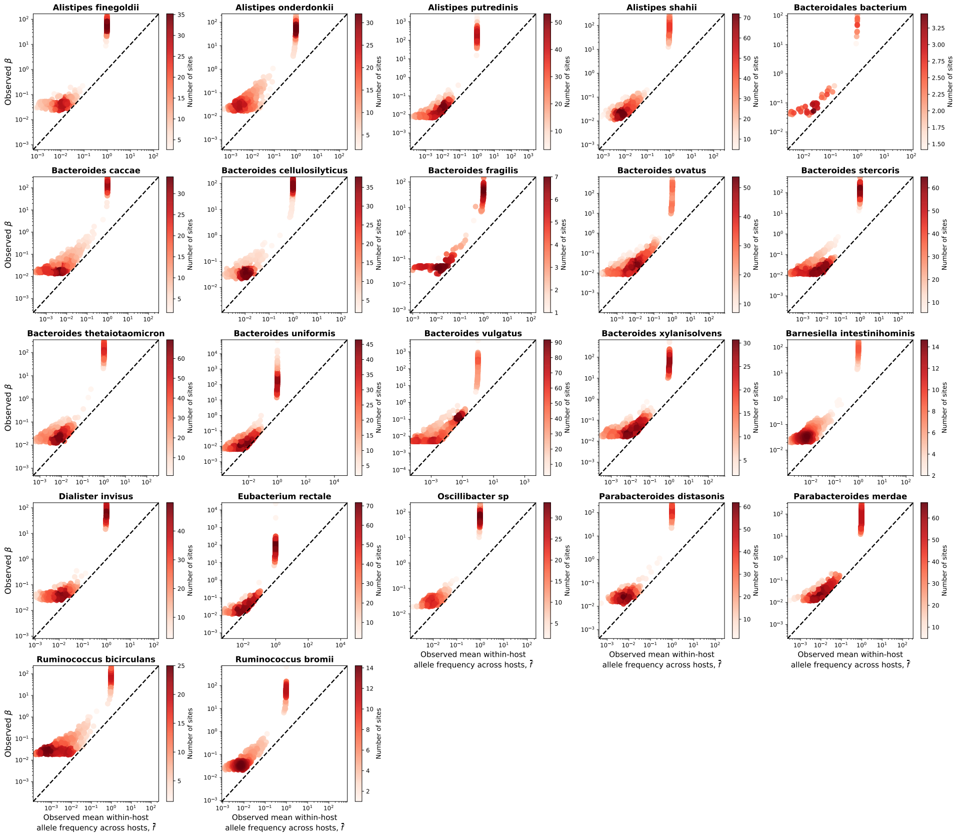


**Figure S13.** The relationship between the empirical estimates of the two parameters of the SLM: the mean allele frequency across hosts ($\bar{f}$) and the squared inverse of the coefficient of variation of frequencies across hosts ($\beta$). Each point is an individual allele. All alleles are at nonsynonymous sites. A total of 1,000 datapoints were sampled without replacement for each subplot. The color of each datapoint is proportionate to the number of sites.
